# Individual differences in responsivity to social rewards: Insights from two eye-tracking tasks

**DOI:** 10.1101/108894

**Authors:** Bhismadev Chakrabarti, Anthony Haffey, Loredana Canzano, Christopher P. Taylor, Eugene McSorley

## Abstract

Humans generally prefer social over nonsocial stimuli from an early age. Reduced preference for social rewards has been observed in individuals with autism spectrum conditions (ASC). This preference has typically been noted in separate tasks that measure orienting toward and engaging with social stimuli. In this experiment, we used two eye-tracking tasks to index both of these aspects of social preference in in 77 typical adults. We used two measures, global effect and preferential looking time. The global effect task measures saccadic deviation toward a social stimulus (related to ‘orienting’), while the preferential looking task records gaze duration bias toward social stimuli (relating to ‘engaging’). Social rewards were found to elicit greater saccadic deviation and greater gaze duration bias, suggesting that they have both greater salience and higher value compared to nonsocial rewards. Trait empathy was positively correlated with the measure of relative value of social rewards, but not with their salience. This study thus elucidates the relationship of empathy with social reward processing.

## Introduction

We are a social species. Humans, from an early stage, generally prefer attending to and interacting with conspecifics compared to objects (Bukach & Peissig, 2009; Legerstee, 1997). This preference for social stimuli has been termed “social preference” and “social motivation” in different theoretical accounts, and is vital for our engagement with the social world (Chevallier, Kohls, Troiani, Brodkin, & Schultz, 2012). A lack of preferential processing of social stimuli can lead to deficits in learning from one’s social environment, and consequent social behavioural deficits in adulthood. One account of conditions marked by deficits in empathy (such as Autism Spectrum Conditions (ASC) and Psychopathy) suggests that empathy deficits seen in these conditions can arise from a core deficit in social reward processing (Dawson, Bernier, & Ring, 2012).

This suggestion is consistent with previous work showing that individuals with high trait empathy have a greater reward-related ventral striatal response to social rewards such as happy faces (Chakrabarti, Bullmore, & Baron-Cohen, 2006). Variations in a key gene expressed within the human reward system (Cannabinoid Receptor gene, *CNR1*) were found to be associated with differences in eye-gaze fixation and neural response to happy faces (but not disgust faces) in three independent samples (Chakrabarti & Baron-Cohen, 2011; Chakrabarti, Kent, Suckling, Bullmore, & Baron-Cohen, 2006; Domschke et al., 2008). In a recent study, Gossen et al. found that individuals with high trait empathy showed greater accumbens activation in response to social rewards compared to individuals with low trait empathy (Gossen et al., 2014).

These studies leave two questions unanswered. First, are rewarding social stimuli preferred when contrasted with an alternative rewarding non-social stimulus? Previous work discussed above has used paradigms where *only* rewarding social stimuli were presented (Chakrabarti & Baron-Cohen, 2011; Chakrabarti, Kent, et al., 2006; Domschke et al., 2008) or where social stimuli were presented on their own (Domschke et al., 2008). Studies that have presented social vs nonsocial stimuli simultaneously, typically have not used stimuli that were rewarding per se (Fletcher-Watson, Leekam, Benson, Frank, & Findlay, 2009; Pierce, Conant, Hazin, Stoner, & Desmond, 2011), but see Sasson & Touchstone, (2014). The importance of adding a rewarding component to social stimuli is that reinforcement signals are important for shaping social behaviour from an early age. If an individual has reduced sensitivity to social reward signals this may lead to atypical social behaviour, as seen in ASC.

Second, does a preference for social rewards manifest through quicker orienting to social rewards, or a longer engagement with social rewards, or both? Both of these phenomena have been observed in infants (Elsabbagh et al., 2013), as well as in young children (Sasson & Touchstone, 2014). It is useful to think of the orienting response as one more related to the salience of the stimuli, while the engagement response as more related to the value of the stimuli. Salience, in this context, refers to motivational salience, or “extrinsic salience”, i.e. a measure of how important a given stimulus is to the observer (Jensen et al., 2007; Zalla & Sperduti, 2013), rather than a stimulus property. On the other hand, ‘value’ of a stimulus refers to how pleasant/unpleasant it is. Salience and value for nonsocial rewards (e.g. food) has been widely studied in primates, and shown to be encoded differently in the brain (Leathers & Olson, 2012; Talmi, Atkinson, & El-Deredy, 2013). However, these processes and individual differences thereof, have not been systematically delineated in the domain of social reward processing in humans.

In this paper, we report two experiments designed to measure two metrics of social reward processing, and relate individual differences in these metrics to trait empathy. The first of these experiments is based on a global effect or centre of gravity effect paradigm (Findlay, 1982; He & Kowler, 1989). In this paradigm, two stimuli are presented peripherally while the participant is asked to make a saccade to a pre-specified target or to simply choose their own target. The saccade tends to deviate (“get pulled”) away from the target toward the other stimulus (Eggert, Sailer, Ditterich, & Straube, 2002; McSorley & van Reekum, 2013). Initial saccades in the global effect task are usually short (~180-230ms) and are known to be influenced more by target salience (Schütz, Trommershäuser, & Gegenfurtner, 2012). This paradigm thus allows for direct attentional competition between social and nonsocial reward targets. The extent to which the saccade gets deviated toward social images compared to nonsocial images can then be used as a metric for relative salience of social rewards.

In the second experiment, we used a preferential looking task, widely utilised in developmental psychology to index preference (Batki, Baron-Cohen, Wheelwright, Connellan, & Ahluwalia, 2000; Fantz, 1958). In a preferential looking task two images are presented side by side and participants are provided with no instructions. This provides an unconstrained setting for participants to fixate wherever they like, and allows them to switch back and forth between the two pictures, for a long duration (usually >=5s). Gaze duration in tasks of this type correlates strongly with self-reported choices and preference ratings (Taylor, Schloss, Palmer, & Franklin, 2013), and has been suggested to encode relative value (Krajbich, Armel, & Rangel, 2010; Shimojo, Simion, Shimojo, & Scheier, 2003). In the current study, we utilised this paradigm to measure preferential gaze duration for social compared to nonsocial pictures, as a putative index of relative value for social compared to nonsocial stimuli.

In order to address the key questions using the paradigms described above, it was necessary to develop a set of stimuli with social and nonsocial rewarding content, that were age-appropriate for an adult sample. A scrambled version of these images were also created to control for the impact of low-level visual features on any comparison between image types (details of image matching parameters are described below in the methods section).

We hypothesised that social rewards will be associated with greater gaze duration bias and saccadic deviation compared to nonsocial rewards, and that trait empathy will be positively correlated with the measures of social preference. We did not have a prior hypothesis on whether empathy will be related to the measures related to salience or value of social rewards, in the absence of prior behavioural data.

## Methods

### Stimuli

40 pairs of positively valenced images were chosen for their social or nonsocial content. Social content was defined as images where one or more humans were visible in the image (e.g. happy couples, babies), while nonsocial content included objects and food items targeted to appeal to a range of individuals (e.g. cupcakes, cars). A subset of these images (15 social, 21 nonsocial) were drawn from the International Affective Picture System (IAPS; Lang, Bradley, & Cuthbert, 1999), while the rest were drawn from publicly available creative common licensed images databases such as Flickr (stimuli set available upon request).

Each social reward image was paired with a specific nonsocial reward image such that they were closely matched on a number of psychological (arousal, valence) and stimulus parameters (contrast, and stimulus saliency. The extent of matching in each of these parameters for each pair of images is depicted in Figure 1. The global RMS and Local RMS contrast were computed as described in (Bex & Makous, 2002). Image saliency (a characteristic of the image calculated based on its low-level visual features) was calculated using the Koch toolbox (Walther & Koch, 2006). Please note, however, that “image saliency” as calculated by the Koch toolbox is a property solely of the image, and is different from our use of the term “salience”, which refers to a property of the image in relation to the observer (how important/relevant an image is to the observer). The confidence intervals for all of these ratios overlap the value of 1, suggesting that there was no significant difference on any of these parameters between social and nonsocial reward images.

**Fig 1.**
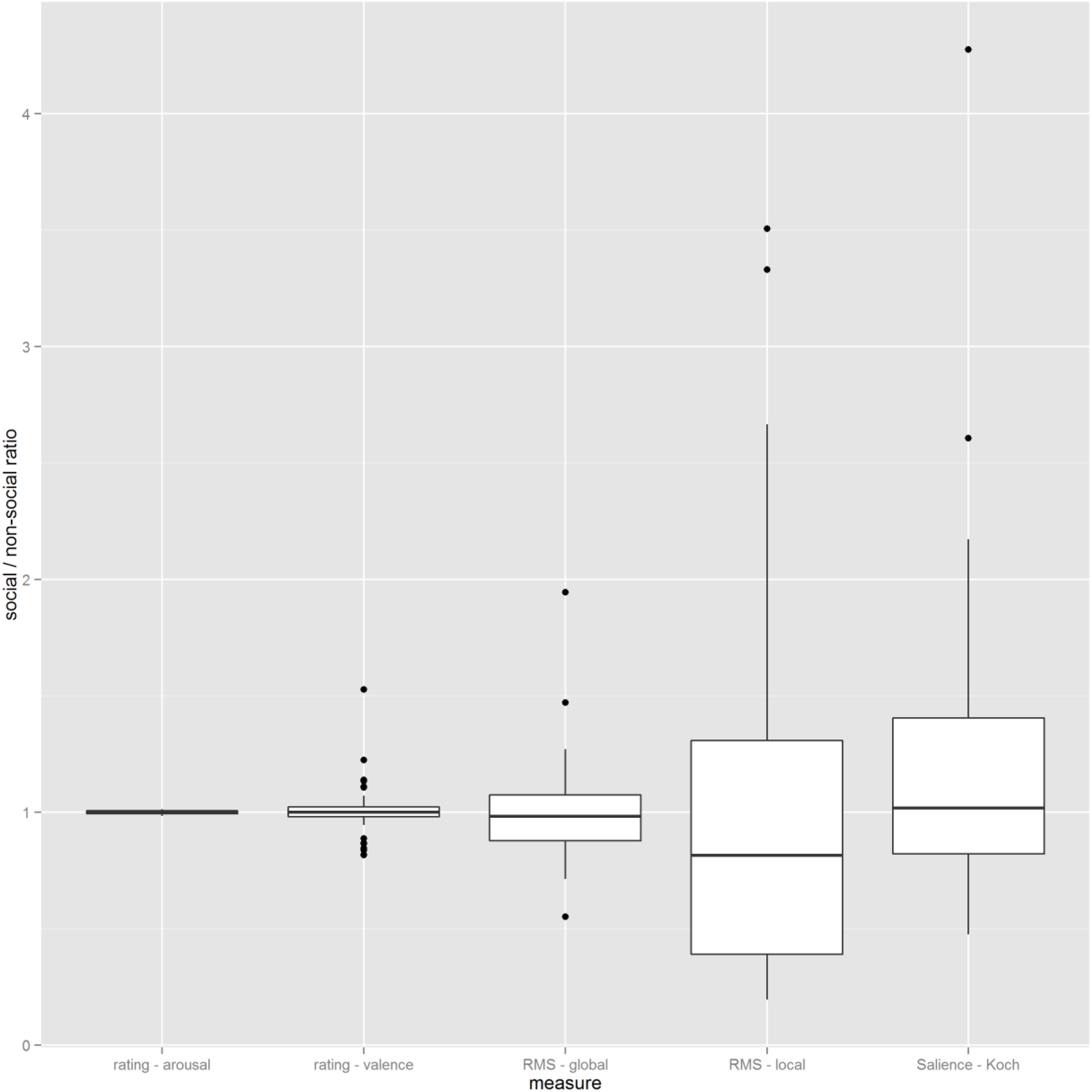
Parameters of matching of social and nonsocial images: The ratio of a number of parameters were calculated for each pair of images (1 social, 1 nonsocial) used as stimuli in both the tasks. These consisted of psychological parameters (arousal and valence ratings) as well as image parameters (Global Root Mean Square[RMS] contrast, local RMS contrast, and stimulus saliency).

In addition, all image pairs were converted into grid-scrambled 10-pixel mosaics to create a control stimuli set, to control for the effect of low level properties of the stimuli such as contrast or colour on any of the measures of interest. All 40 image pairs were used in both tasks described below.

### Tasks

#### Global effect (GE) task

The GE task was based on a modified version of the original task that has been used in studying response to emotional stimuli (McSorley & van Reekum, 2013). The stimuli were presented as shown in Figure 2. Each trial began with the presentation of a central fixation cross. After 800-1200 ms this disappeared and immediately reappeared 6 degree of visual angle to the left or right of centre on the horizontal meridian. Participants were instructed to look at wherever the fixation cross reappeared. This required making a straight direct saccade from the centre to the periphery of the screen.

**Fig 2.**
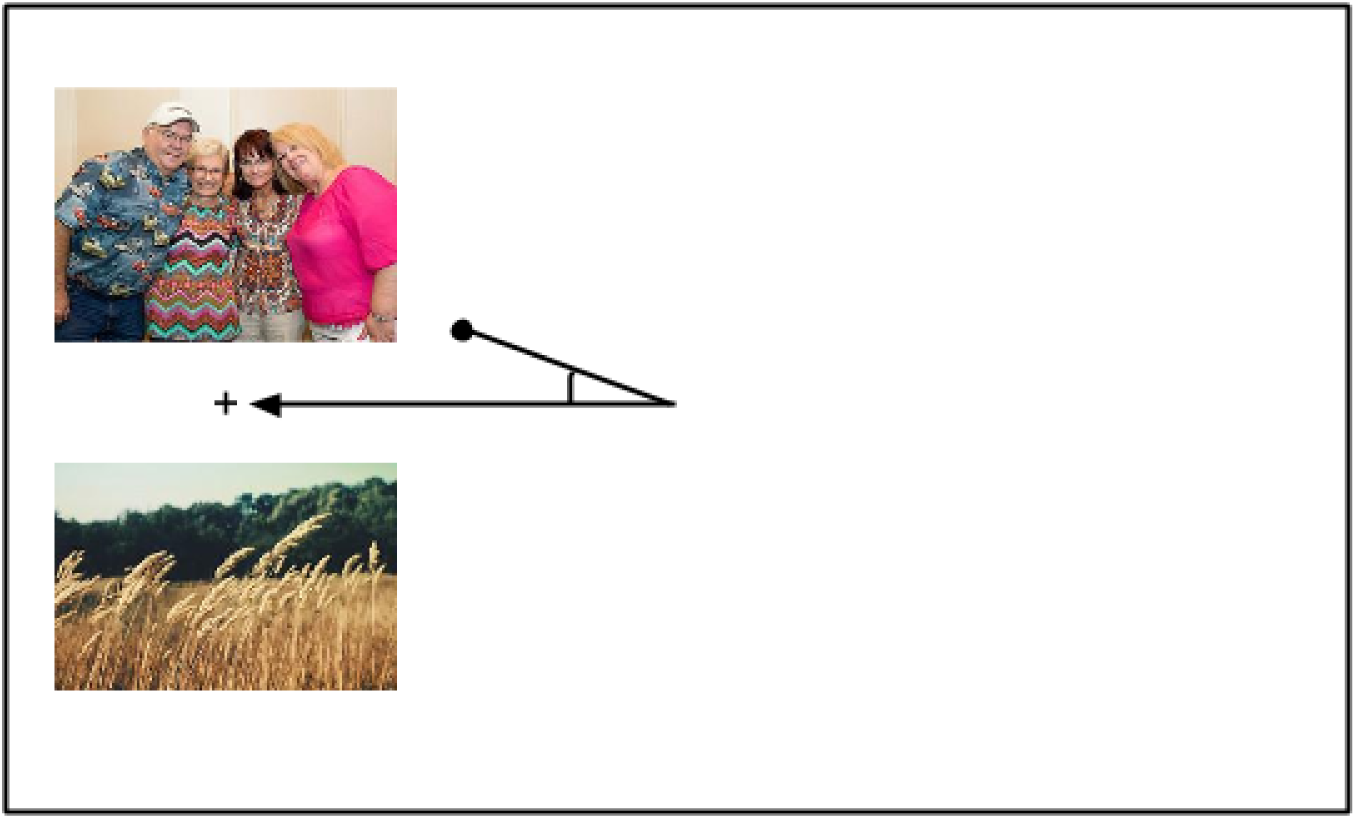
The layout of a trial in the Global Effect task in which the images appear to the left of the initial fixation cross. The angle of deviation towards the top or bottom image is calculated as the difference between the participant’s first saccade (represented by a black circle) and the shortest path between the initial location of the fixation cross and the location where it reappears (on the left or right of the screen). Social and nonsocial reward images appeared in pairs, and were presented to the right or the left of the initial fixation cross for an equal number of times. Participants were instructed to look only at the fixation cross, and ignore the images. The images were 5.59° wide and 4.19° tall. There was a .28° distance between the reappeared fixation cross and the presented pictures. The distance between the initial and the reappeared fixation is 6°. The images in this figure were not used in the study; public domain images have been inserted due to copyright restrictions. The full set of stimuli is available on request from the first author.

Each of the 40 pairs of (social and nonsocial) images was presented in its scrambled and unscrambled form, thus comprising 80 trials. Stimulus type (scrambled/unscrambled, social/nonsocial) and spatial location (left/right, top/bottom) were counterbalanced across participants.

#### Freeview (Preferential Looking) task

The stimuli were presented as shown in Figure 3. The 40 pairs of images (social and nonsocial) and 40 control pairs of scrambled images were presented in a pseudorandom sequence side by side (see Figure 3). Each trial began with the presentation of a central fixation cross (“+” 0.28 degree of visual angle). Once participants fixated on the cross an automated drift correct procedure was performed using four head cameras that corrected for any slight movements of the participants. Following this, the fixation cross was removed and a pair of social and nonsocial images (each image 5.59 deg x 4.19 deg) were immediately presented for 5 seconds to the left and right. During the trial the participant was free to look wherever they chose. This was followed by a 1500ms intertrial interval.

**Fig 3.**
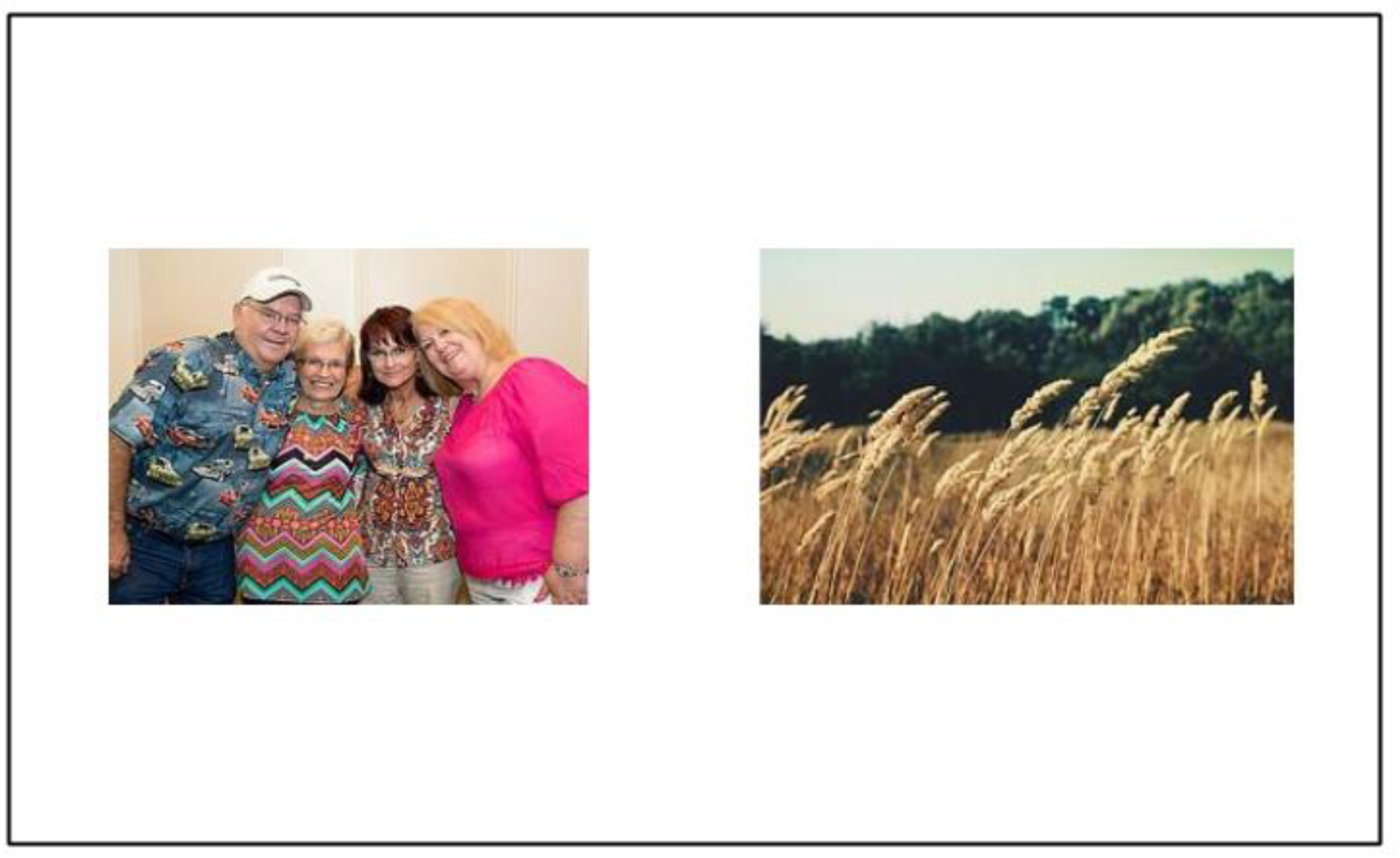
The layout for a trial in the Freeview task in which participants were free to look at paired social and nonsocial images that were presented side by side. The average duration participants spent looking at the social vs. nonsocial images was measured. Images were presented 4.46° apart, and were 5.59° wide and 4.19° tall. Images were presented for 5000ms. The images in this figure are presented solely for representative purpose, these were not used in the study. The full set of stimuli is available on request.

#### Procedure

After giving informed consent participants were briefed about both tasks. Head movements were constrained with a chin-rest, which held participants so their eyes were in-line with the horizontal meridian of the screen, at a viewing distance of 1m. The eye-tracker was calibrated using a standard 9 point grid, carried out at the beginning of the experiment. Calibration was only accepted once there was an overall difference of less than 0.5 degree between the initial calibration and a validation retest: in the event of a failure to validate, calibration was repeated. The order of the two tasks was counterbalanced across participants.

Eye movements were recorded using a head-mounted, video-based, eye-tracker with a sampling rate of 500 Hz (Eyelink II, SR Research). Viewing of the display was binocular and we recorded monocularly from observers’ right eyes. Stimuli were presented in greyscale on a 21” colour monitor with a refresh rate of 75 Hz (DiamondPro, Sony) using Experiment Builder (SR Research Ltd.). Participants completed the Empathy Quotient (EQ; Baron-Cohen & Wheelwright, 2004) questionnaire online.

#### Participants

77 participants (42 females; mean age= 21 years, 1 month, s.d. = 3 years and 5 months) drawn from in and around the University of Reading campus completed the FV task. One participant’s data in the FV task was removed because gaze data was captured on less than 75% of scrambled trials. 74 of the FV participants completed the GE task. GE data for 7 participants were discarded due to capturing fixations on fewer than 75% of scrambled or unscrambled trials. All participants had normal or corrected to normal vision. Of the remaining participants, 68 FV (38 female) and 61 GE (33 female) participants completed the online EQ questionnaire. The study was approved by the University of Reading Research Ethics Committee.

#### Data analysis

The data was analysed and figures were generated with R using the ggplot2 (Wickham, 2009) and grid (Team, 2012). The global effect in response to social stimuli was measured as the average deviation towards social vs nonsocial images. During each trial the angle of the first saccade identified was calculated relative to how far it was off the line between where the fixation cross initially appears and where it reappears, and whether it was toward the social or nonsocial image (See Figure 2). A positive average deviation indicates a bias towards social stimuli whereas a negative value indicates a bias towards nonsocial stimuli. Average deviation was calculated separately for scrambled and unscrambled images.

Preferential looking for social stimuli in the FV task was measured as the proportion of gaze duration (dwell time) on social images on each trial.

All test statistics presented in the following section are 2-tailed.

## Results

### Global effect task

For unscrambled images, a one sample t-test against a test value of 0 (which corresponds to no significant deviation toward social/nonsocial image) found that *average deviation* was significantly more toward social than nonsocial images (t (66) = 8.409, p<.001, Cohen’s d = 1.027). This was not true for scrambled images (t (66) = -.392, p = .697, Cohen’s d =.048; see Figure 4a). A direct comparison of the extent of deviation toward social images in the unscrambled and the scrambled conditions revealed a significant difference in *average deviation* (t(66) = 7.371, p <.001; Cohen’s d = 1.253; see Figure 4a).

**Fig 4.**
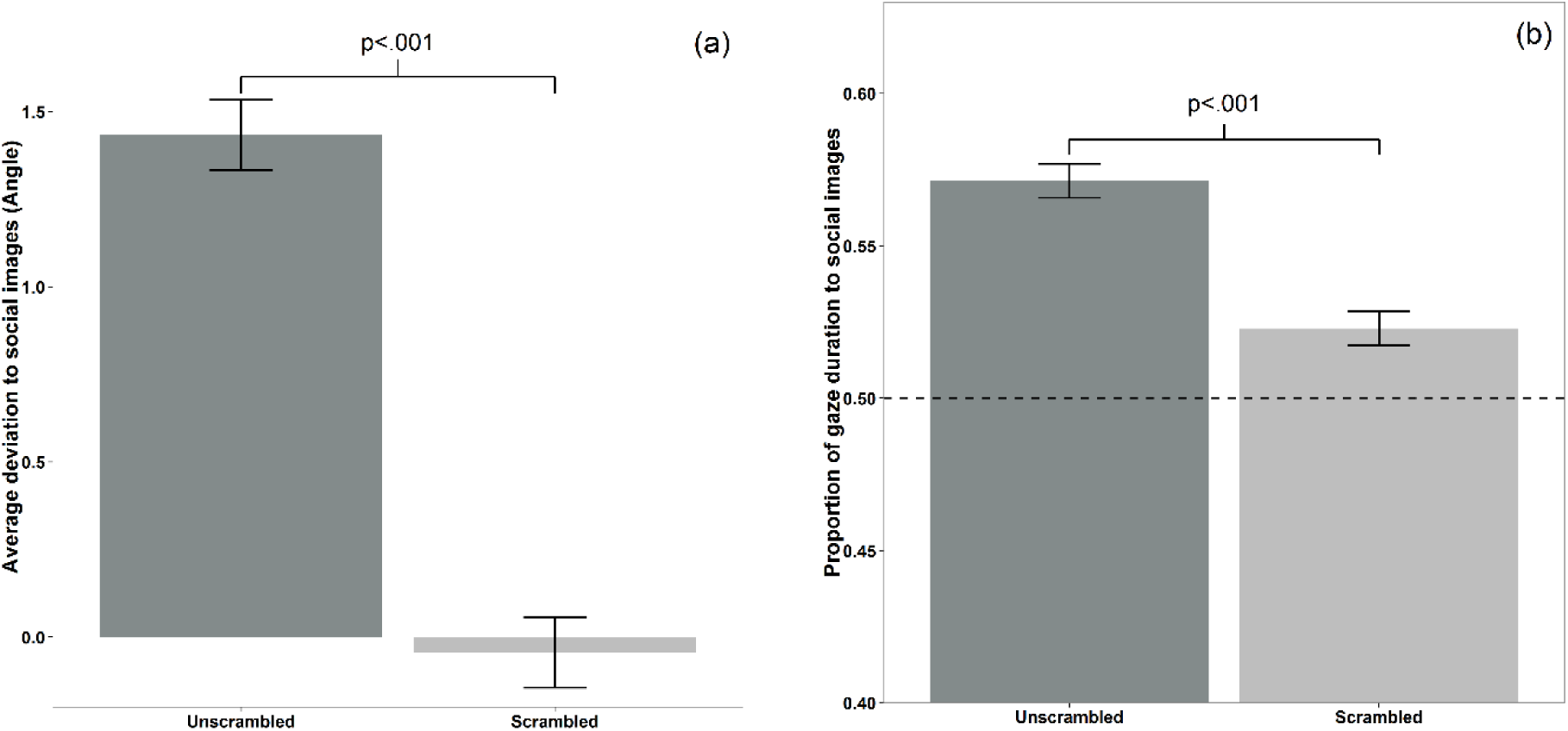
(a) Bar graph showing the average angle of deviation towards social images during the Global Effect task. There was a significant sociality bias for *average deviation* on unscrambled images (p<.001), but not scrambled images (p=.697). The error bars reflect within- subject errors, calculated using the Cousineau (2005) method. (b) Bar graph of the proportion of duration looking to social images during the Freeviewing task. Gaze duration was significantly longer to social images for unscrambled (p<.001) and scrambled images (p<.001). Values above the dotted line at .5 indicate a bias to social images.

#### Correlations

There was no correlation between EQ and the *average deviation* toward social reward images for unscrambled (r (59) =.059, p =.652) or scrambled images (r (59) = -.114, p =.382).

### Freeview task

#### Main effects

One sample t-tests against a test-value of .5 (corresponding to equal proportion of dwell time to social and nonsocial images) found that the proportion of gaze duration to social images was significantly higher for both unscrambled images (t (75) = 6.664, p<.001, Cohen’s d =.764) and scrambled images (t (75) = 4.362, p<.001, Cohen’s d =.5; see Figure 4b). A paired samples t-test found that proportion of gaze duration to social images was greater for unscrambled images than scrambled images (t (75) = 4.285, p<.001, Cohen’s d =.652; see Figure 4b).

#### Correlations

There was a positive correlation between EQ and proportion of gaze duration to social images in unscrambled images (r(66) =.278, p=. 022), but no significant correlation between EQ and proportion of gaze duration to social images in scrambled images (r(66) = -.198, p =.106; see Figure 5). To directly test the difference between these two dependent correlations Steiger’s Z was calculated (Steiger, 1980). This showed a significant difference between the correlations (Steiger’s Z = 2.95, p =.003).

**Fig 5.**
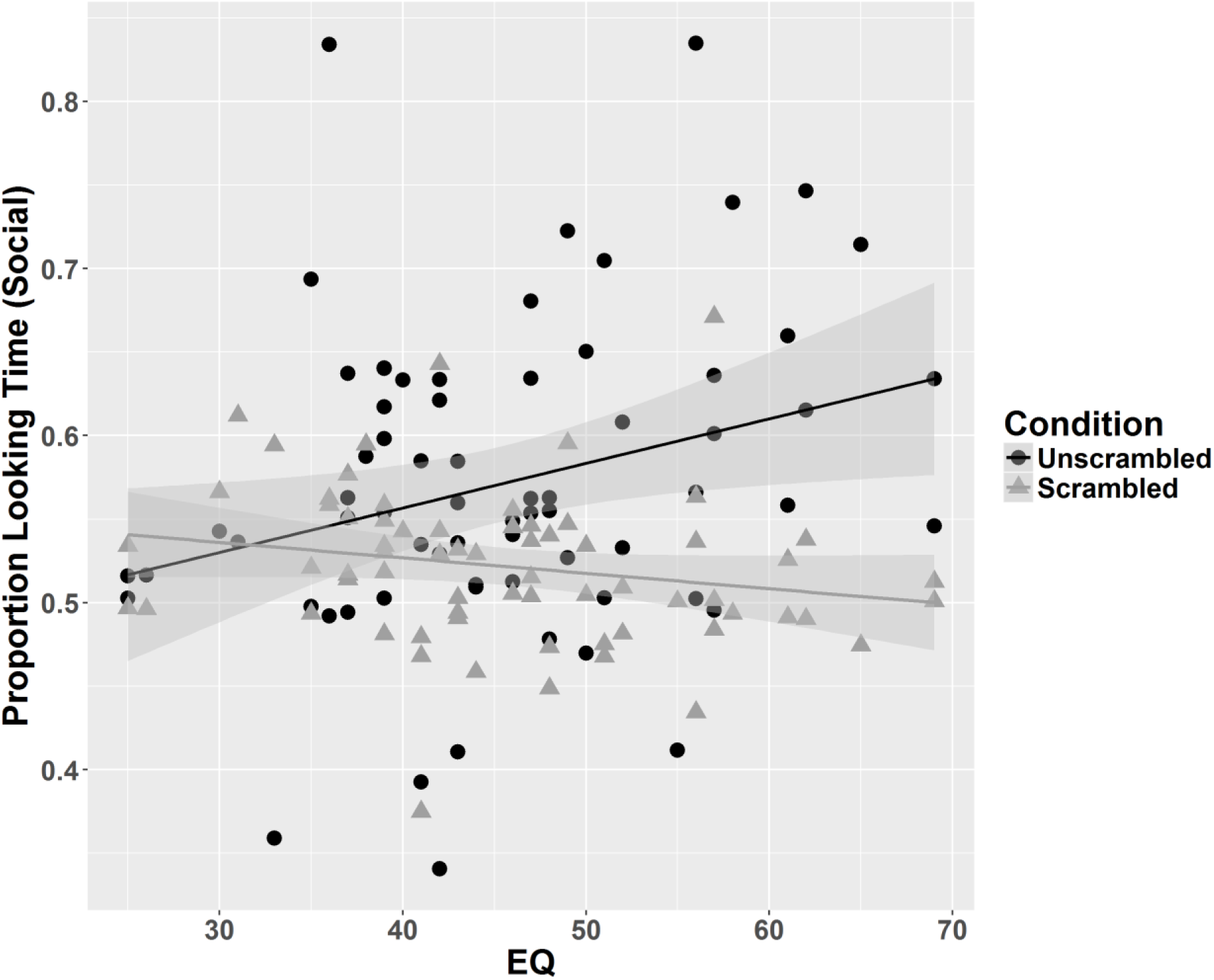
Scatter plot of empathy quotient (EQ) scores with the proportion of gaze duration to social images for unscrambled images (black line and circles) and scrambled images (grey line and triangles). Greater empathy predicts more time spent looking at social images than nonsocial images when they are unscrambled (r(66) =.277, p=. 022) but not scrambled (r(66) = -.198, p =.106).

## Discussion

In this study we developed a new set of ecologically valid stimuli depicting of social and nonsocial rewards. Using two separate tasks with this stimulus set in the general population, we found that social reward images evoked greater saccadic deviation and preferential looking than nonsocial reward images, after having controlled for differences in low-level visual properties. This result supports similar work on dynamic scenes which showed that low-level stimulus features are not able to explain the gaze response to social stimuli (Coutrot & Guyader, 2014). Importantly, the preferential gaze bias toward social reward images was proportional to individual differences in trait empathy. This relationship with empathy was seen only with the measure of engagement (i.e. the preferential gaze experiment), and not for the measure of orientation (i.e. the experiment measuring saccadic deviation).

Humans orient quickly to social stimuli from an early age, across sensory modalities (Dawson, Meltzoff, Osterling, Rinaldi, & Brown, 1998; Johnson, Dziurawiec, Ellis, & Morton, 1991; Mosconi, Steven Reznick, Mesibov, & Piven, 2009). Typically developing children orient more to social stimuli, when these are presented within an array of nonsocial stimuli (Sasson, Turner-Brown, Holtzclaw, Lam, & Bodfish, 2008). Orienting responses are driven primarily by the salience of the target, and hence these results suggest that social stimuli are generally regarded as more salient than nonsocial stimuli. However, none of these paradigms have measured the global effect which relies on quick saccades that are influenced more by target salience than value (Schütz et al., 2012). These results are therefore consistent with the literature on infants and young children, and points to a higher salience of social compared to nonsocial reward images in adults.

Relative gaze duration in paradigms where two stimuli are presented simultaneously may index the relative value of the two targets and two computational models have been proposed (gaze cascade and drift diffusion models) to relate gaze duration bias and the relative value of targets (Krajbich et al., 2010; Shimojo et al., 2003). Preferential looking paradigms have been used widely in developmental psychology, where it has been shown that infants look longer at social compared to nonsocial stimuli (Johnson et al., 1991; Jones & Klin, 2013; Simion, Regolin, & Bulf, 2008). This suggests that in typically developing infants social stimuli in general have a higher value than nonsocial stimuli. However, these stimuli are usually not matched for visual properties. Our results are consistent with these results, and show that social rewards may have a higher value than nonsocial rewards. Importantly, this difference was not driven by a difference in stimulus arousal/valence, or by differences in low-level properties of the images that we tested.

Empathy was found to be directly proportional to the gaze duration for social compared to nonsocial reward images. This suggests that highly empathic individuals may attribute a greater value to social rewards compared to nonsocial rewards. This is consistent with experiments which found greater striatal activation to happy faces in individuals with high trait empathy (Chakrabarti, Bullmore, et al., 2006). Janowski et al. (2013) showed that empathic choice is influenced by processing of value in the medial prefrontal cortex in a choice task. Jones & Klin (2013) showed that infants who go on to develop autism show a progressively reduced gaze fixation toward faces when presented simultaneously with objects in a naturalistic video. Individuals with ASC score low in questionnaire measures of empathy (Baron-Cohen & Wheelwright, 2004). These converging lines of evidence suggest that individuals low in empathy might show reduced gaze duration for social vs nonsocial stimuli, a suggestion supported by our results. Future work should explore if these effects are magnified or reduced, if nonsocial reward images are chosen to be of high interest to individual participants, similar to the approach taken by Sasson and colleagues (Sasson & Touchstone, 2014; Sasson, et al., 2011). Another potential avenue for future exploration would be to test if such indices of social preference hold true if stimuli of neutral valence are used.

### Conclusion

In this set of two experiments we used a new set of images of social and nonsocial rewards and showed that social rewards are associated with greater saccadic deviation and higher gaze duration compared to nonsocial rewards. This social advantage persists even after minimising differences in arousal/valence of these images and low level visual properties. We found that trait empathy was correlated positively to the gaze duration bias for social rewards, but not with the saccadic deviation toward social rewards. This results points to a potential distinction between two important aspects of social reward processing (i.e. salience and value), and clarifies their relationship with phenotypic dimensions relevant to ASC. Future research should directly test these different parameters of social reward processing in individuals with atypical empathy profiles, such as those with ASC and Psychopathy.

## Acknowledgements

The authors acknowledge Natalie Kkeli, Violetta Mandreka, Charlotte Whiteford and Kara Dennis for help with data collection. BC is supported by a MRC New Investigator grant, CPT was supported by the University of Reading Research Endowment Trust Fund, ATH is supported by an ESRC-MRC interdisciplinary doctoral studentship, LC is supported by Sapienza University of Rome studentship.

